# Droplet-digital Cas13a assay enables direct single-molecule microRNA quantification

**DOI:** 10.1101/748939

**Authors:** Tian Tian, Bowen Shu, Lei Liu, Xiaoming Zhou

## Abstract

The direct quantification of microRNA at the single-molecule level is still challenging. Herein, we developed a droplet-digital Cas13a assay (ddCA) as a general approach for the amplification-free and absolute quantification of single unlabeled miRNA molecules with single-nucleotide specificity. We demonstrate its simplicity, precise quantification capability, excellent specificity and broad applicability by analyzing microRNAs from synthetic and cell lines-derived materials.

---

Insights into the roles of microRNAs (miRNAs) as key regulators in a range of biological processes, promising biomarkers for disease diagnostics, and novel targets for therapeutics have attracted increasing attention to miRNAs recently^1–3^. The accurate quantification of miRNAs is critical to better understanding these roles. However, many properties unique to miRNAs pose challenges for their precise quantification, such as short lengths, sequence similarity, low abundances, and the wide range of expression variance^4^. Most current methods used for highly sensitive miRNA detection require lengthy sample preparation or amplification steps that can bias the results^5^. Approaches capable of direct detection of miRNA improving the quantitation ability but often suffer from insufficient sensitivity and specificity; or require the complicated labelling and chemical modification of the target (see **Supplementary Note 1** and **ref**.^1–17^).

To overcome these challenges, we developed droplet digital a Cas13a assay (ddCA) for the direct quantification of miRNAs at the single-molecule level. A key unique feature of this approach is the use of Cas13a^6–9^ to provide specific RNA recognition and signal amplification within picoliter-sized droplets, thus enabling direct counting of the number of RNA targets from the resulting droplets (**Fig.1a**). This enables the digital quantification of miRNAs without reverse transcription or nucleic acid amplification. Due to its simplicity, the entire workflow can be conducted by coupling the use of a syringe-vacuum driven microfluidic droplet chip with standard fluorescence microscopy (**Fig.1b**, **Online Methods**, **Supplementary Fig.S1** and **Video S1**). *Via* a simple operational procedure, highly uniform microdroplets (30 μm in diameter with a coefficient of variation below 1.7 %) are robustly generated, and subsequently self-assembled into a droplet array for imaging-based 2D droplet counting. (**Fig.1c**, **Supplementary Fig.S2**).

**Figure 1.**
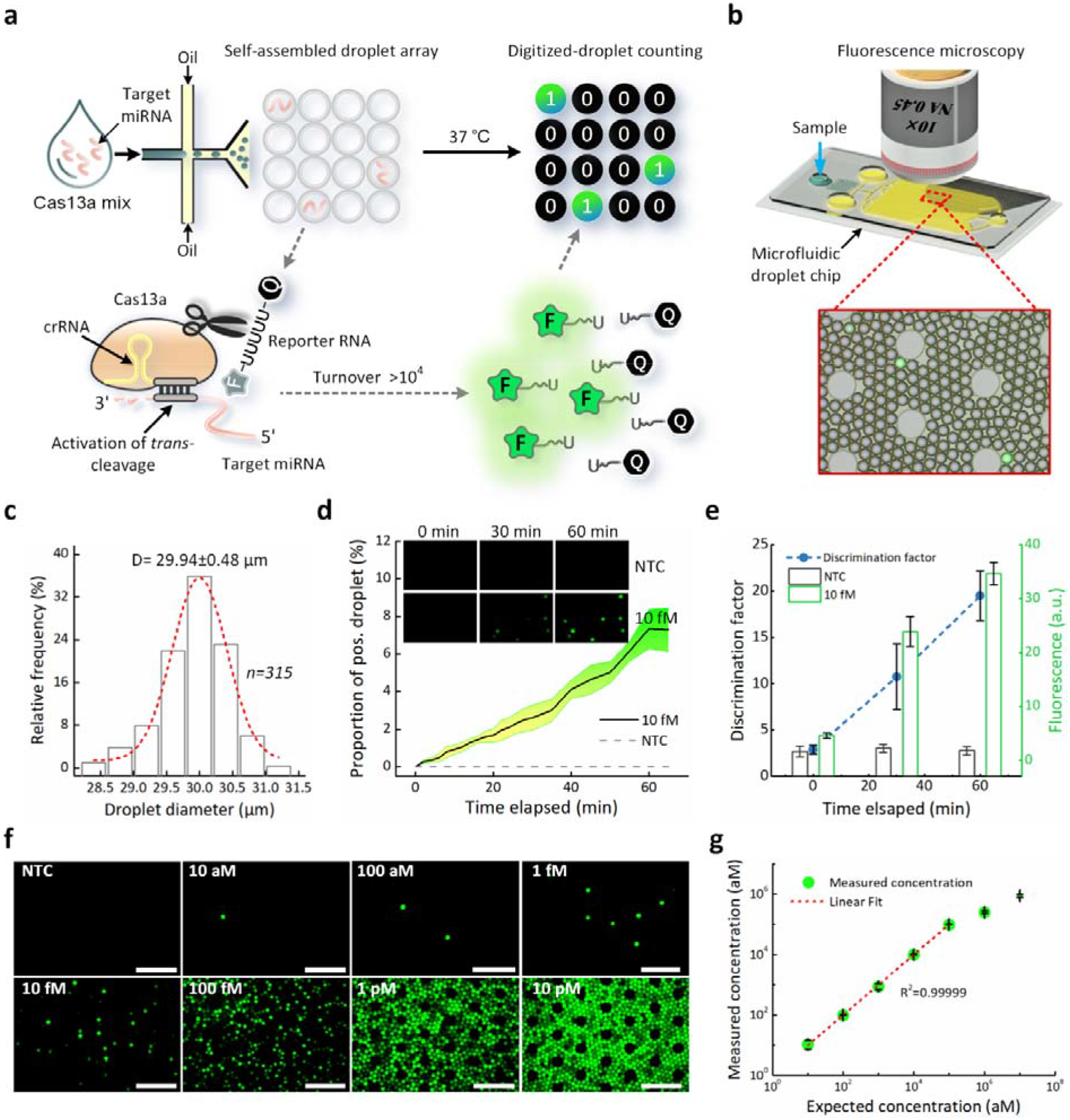
Single-molecule miRNA quantification using ddCA. **(a)** Schematic of ddCA. MiRNAs (pale red) together with a Cas13 mix are emulsified with oil into thousands of picoliter-sized droplets, each containing approximately 0 to 1 molecules of target miRNA. Once recognized by a crRNA (yellow), one target miRNA will induce cleavage of 10^4^ quenched fluorescent RNA reporters (F, fluorescence; Q, quencher), thus yielding a fluorescently positive droplet. **(b)** Workflow of ddCA. By coupling a microfluidic droplet chip with fluorescence microscopy, highly uniform microdroplets are robustly generated and self-assembled into a high-density monolayer droplet array for fluorescence imaging-based 2D droplet counting. **(c)** Size distribution of the droplets. **(d)** Time course showing the proportion of positive droplets resulting from single-molecule miRNA detection during 37 °C incubation. **(e)** The background-subtracted fluorescence difference between the positive and negative droplets during single-molecule miRNA detection *via* ddCA (mean ± SD). **(f)** Representative end-point fluorescence images of various miRNA-17 concentrations obtained *via* ddCA (10× magnification, scale bar=200 μm). **(g)** Quantification range of ddCA (mean ± s.e.m, n = 3).

To examine the feasibility of single-molecule miRNA detection, we used 10^4^ aM (~6×10^3^ copies per μL) synthetic miRNA-17 as input, which permits ≤ 1 molecule contained in a droplet with a probability of 99.67%. We determined the time course of the proportion of positive droplets (*PPD*) during in-situ isothermal incubation. We found that the *PPD* reached a plateau within 60 minutes, with a measured value (7.3%) that agreed closely with the predicted value (8.4%) (**Fig.1d**). Meanwhile, an evident difference (≈20-fold) in the fluorescence signal between the positive and negative droplets could be measured, which was favorable for accurate digitization (**Fig.1e**). Thus, a 60-min single-temperature incubation is sufficient for single-molecule miRNA detection *via* ddCA.

To investigate the quantification capability of ddCA, we detected serial dilutions of known concentrations of synthetic miRNA-17 from 10^1^ to 10^7^ aM. Intuitively, the end-point fluorescence images showed the consistency between the concentration and the *PPD* (**Fig.1f**). Furthermore, we observed an excellent (R^2^>0.9999) linear response between the measured concentration and the input concentration in the dynamic range from 10^1^ aM to 10^5^ aM (**Fig.1g**). Here, the lower detection limit of ddCA was determined to 3 aM (~ 2 copies per μL) with a 95% confidence interval, when considering that the imaging chamber of our microfluidic chip is designed to accommodate at least 10^5^ droplets. We also noticed that the measured concentration was lower than the input concentration when the target concentration was over 10^5^ aM, despite using *Poisson* distribution correction. Nevertheless, the linear dynamic range spanned 5 orders of magnitude, and along with the single-molecule level sensitivity, this allowed for the precise determination of the profile of most miRNA expression variation^5^.

Notably, superior performance could be achieved within the microdroplets compared to bulk volumes using the Cas13a assay (**Fig.2a** and **Supplementary Fig.S3**). The reduced reaction volume and the droplet-based digital assay method might be key factors contributing to the superior performance of the assay (specifically detailed in the **Supplementary Note 3**). In the case of the presence of a single miRNA molecule (**Fig.2b**), the volume reduction from 25 μL to 14 pL will increase the effective target concentration over 10^6^ times; thus, it is not surprising that the Cas13a assay in the microdroplet attained significant sensitivity enhancement. In contrast to well-established methods (**Fig.2c-d**), our amplification-free approach has an extended lower limit of quantification and a dynamic range matching that of qRT-PCR assay but avoids the risk of false positives due to carryover contamination and the need for internal reference or calibration. Furthermore, a larger dynamic range and an equal sensitivity were obtained by ddCA compared to ddPCR, presumably due to the benefits from the larger number of droplets applied and the avoidance of droplet loss during droplet transfer and thermocycling^10–11^.

**Figure 2:**
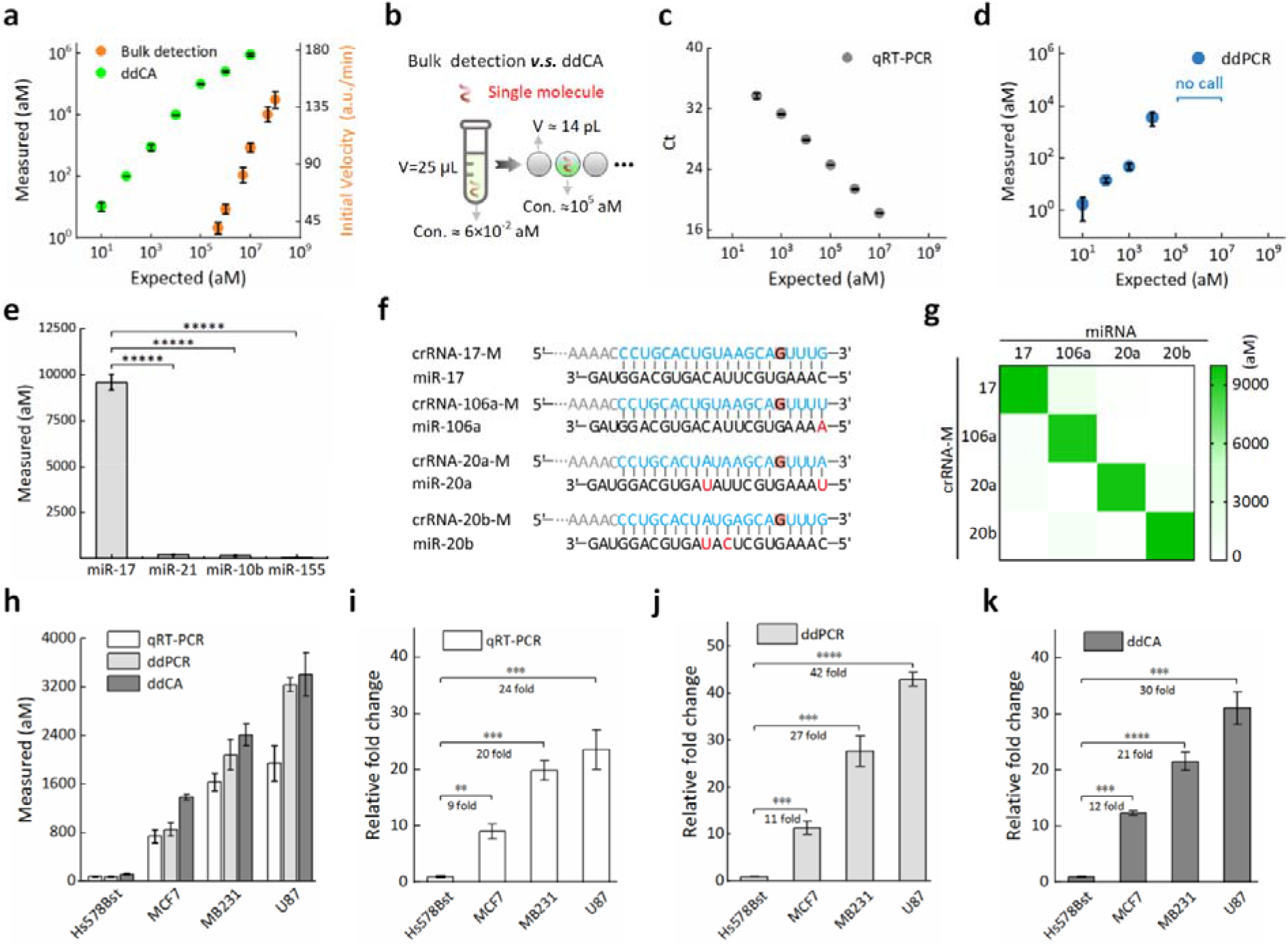
Assay performance of ddCA. **(a)** Quantitation of synthetic miRNA-17 using ddCA and bulk fluorescence detection. **(b)** Schematic showing the concepts of bulk detection and ddCA. The detection of single-molecule miRNAs in microdroplets substantially increases the local concentrations of the target miRNAs. **(c-d)** Quantitation of synthetic miRNA-17 using qRT-PCR and ddPCR. **(e)** ddCA assay for detecting four microRNAs from different families utilizing crRNA complementary to miR-17 (two-tailed Student’s *t* test; *****, p < 0.00001). **(f)** Sequences of crRNAs with synthetic mismatches (red highlight) complementary to four homologous miRNAs: miR-17, −106a, −20a, and −20b. SNPs in the target are colored in red. **(g)** Heat map analysis of the orthogonal identification of four homologous miRNAs using ddCA. **(h)** Quantification of miR-17 expression in the total small RNA extracts (0.1 ng) using qRT-PCR, ddPCR and ddCA. **(i-k)** Relative expression of miR-17 by qRT-PCR, ddPCR and ddCA (two-tailed Student’s *t* test; ***, p < 0.001; ****, p < 0.0001). All the experiments were repeated at least 3 times, and the bars represent the mean ± s.e.m.

To check the specificity of ddCA, we designed a truncated crRNA with a 20-nt spacer to discriminate four miRNAs from different families including miRNA-17, -10b, -21, and -155 (**Supplementary Fig. S4**). MiRNA-17 complementary to the truncated crRNA yielded over a 50-200-fold increase in outcomes compared to those obtained with the other three miRNAs when using an equal amount of input miRNA, exhibiting a significant difference (*p*<10^−5^) in selectivity owing to sequence heterogeneity (**Fig.2e**). Cas13a has tolerance for one mismatch but is sensitive to two or more mismatches in the crRNA spacer: target duplex^12–13^. To circumvent this issue, we introduced an artificial mismatch in the crRNA spacer sequences to further enhance specificity (**Fig.2f**). The orthogonal testing results demonstrated that our ddCA successfully discriminated the homologous microRNAs (−17, −106a, −20a, and −20b) which belong to the same family and differ by just one or two nucleotides (**Fig.2g** and **Supplementary Fig. S5**). These results indicated that ddCA has excellent specificity for discriminating similar miRNA sequences, even those with single nucleotide differences.

To assess the potential utility for precise miRNA profiling, we used ddCA to quantify the miR-17 expression level in four different cell lines, including human breast normal cells (Hs578Bst)/adenocarcinoma cells (MCF-7 and MDA-MB-231), and human glioma cells (U87). Overall, compared with the results from qRT-PCR and ddPCR, a slightly higher level of miRNA-17 expression was measured in each cell line using ddCA (**Fig. 2h**). This might be attributed to the high fidelity of our amplification-free quantification, which avoids sample loss during reverse transcription and amplification bias. Meanwhile, we observed similar variations in the miRNA-17 relative expression levels in different cell lines using the aforementioned three methods, and all these methods revealed significant differences in miRNA-17 expression between the normal cell lines and the cancer-related cell lines (**Fig.2i-k**). This result suggests that our ddCA method enables the precise quantification of miRNA expression levels in a high nucleic acid background.

In conclusion, we presented a simple, high-fidelity, direct detection method for the absolute quantification of miRNA at the single-molecule level. The ddCA method provides a unique combination of performance parameters that outperforms those of other state-of-the-art miRNA detection technologies (see **Supplementary Note 1**), so we envision that this approach may serve as a next-generation platform for RNA quantitation and will find broad application in both basic research and clinical diagnostics.

## Methods

### LbuCas13a protein expression and purification

Briefly, *Escherichia coli* Rosetta (DE3) cells harbouring pET28a-His_6_-SUMO-LbuCas13a plasmids, a gift from professor Yanli Wang, were cultured at 37 °C until OD_600_=0.6, using Terrific Borth growth media (OXOID) with 100 mg/mL ampicillin (Sangon). Then, 100 μM isopropyl-1-thio-b-D-galactopyranoside (IPTG, sigma) was added to the culture to induce protein expression at 26 °C for 4 h. The cells were harvested and lysed by sonication in lysis buffer (20 mM Tris-HCl, 1 M NaCl, 10% glycerol, pH 7.5). The cell lysate was separated by centrifugation at 8,000 r.p.m. at 4 °C, and the collected supernatant was incubated by Ni-NTA agarose (Abiotech, Jinan, China). The bound protein was washed with washing buffer (20 mM Tris-HCl, 150 mM NaCl, 20 mM imidazole, 10% glycerol, pH 7.5) and eluted into an elution buffer (20 mM Tris-HCl, 150 mM NaCl, 500 mM imidazole, 10% glycerol, pH 7.5). Then, the His6-SUMO tags was digested using Ulp1 protease. After that, the resulting protein was purified using Heparin agarose (Abiotech, Jinan, China). The bound protein was washed using washing buffer (20 mM Tris-HCl, 150 mM NaCl, 10% glycerol, pH 7.5) and eluted into elution buffer (20 mM Tris-HCl, 1 M NaCl, 10% glycerol, pH 7.5). The final protein was dissolved in protein storage buffer (20 mM Tris-HCl, 1 M NaCl, 50% glycerol, pH 7.5) and stored at −80 °C for use. (**Supplementary Fig. S6**)

### *In vitro* crRNA transcription

DNA templates containing T7 promoter for *in vitro* crRNA transcription were prepared with the complementary sequence, *via* gradient annealing (95 °C for 5 min followed by cooled to room temperature at a ramp rate of 5 °C/min). T7 transcription reaction, including DNA templates, T7 RNA polymerase (NEB), NTPs mix in 1× RNAPol reaction buffer (40 mM Tris-HCl, 2 mM spermidine, 1 mM DTT, 6 mM MgCl_2_, pH 7.9), was performed at 37 °C for 4 h. The concentration of crRNA was measured by Nanodrop 2000 (Thermo Fisher Scientific). The resulting crRNA was validated by PAGE and then stored at −80°C until use. (**Supplementary Fig. S7**)

### Cell culture and cell total small RNA extraction

Hs578Bst, MB231, MCF7, U87 cell lines were grown in Dulbecco’s modified Eagle medium (Gibco) and supplemented with 10% FBS (EXCELL BIO) at 37 °C in a humid atmosphere containing 5% CO_2_. Total small RNA was extracted from cell lines mentioned above using RNAiso for small RNA (Takara) according to the manufacturer’s protocol.

### Bulk fluorescence detection

Unless otherwise indicated, 25-μL Cas13a mix was prepared with 20 nM LbuCas13a, 10 nM crRNA, 500 nM FQ5U RNA reporter, a certain amount of target microRNA, and 1× reaction buffer (10 mM Tris-HCl, 1.5 mM MgCl_2_, 50 mM KCl, pH 8.9). The bulk fluorescence detection was performed at 37 °C for 40 min using Thermal Cycler Dice Real Time System (Takara), with fluorescence was collected every minute.

### Quantitative real-time PCR experiment

The reverse transcription (RT) cocktail contains 100 nM stem-loop RT primer, dNTPs mix (0.5 mM each) (Takara), synthetic miRNA-17 or 100 ng extracted small RNA, 1 unit/μL RNase inhibitor (NEB), 5 unit /μL SMART MMLV reverse transcriptase (Takara), and 1×MMLV buffer. The RT reaction was carried out at 42 °C for 60 min, then terminated by heating at 70 °C for 15 min. For real-time PCR reaction, 50 μL reaction mixture consisted of 2 μL cDNA, 800 nM forward primer, 400 nM reverse primer, 250 nM Taqman^®^ MGB probe, and 1×Probe qPCR Mix (Takara) was performed by Thermal Cycler Dice Real Time System (Takara). Cycling conditions: 95 °C × 30 s (1 cycle), 40 cycles of 95 °C × 5 s and 60 °C × 1 min.

### Droplet digital PCR experiment

For ddPCR reaction, 25 μL reaction mixture comprised 1 μL cDNA, 800 nM forward primer, 400 nM reverse primer, 250 nM Taqman^®^ MGB probe, and 1×ddPCR supermix for probes (BIO-RAD) was performed by QX200 Droplet Digital PCR system (BIO-RAD). Cycling conditions for ddPCR reaction is 95 °C × 10 min, 40 cycles of 94 °C × 30 s and 60 °C × 1 min, then 98 °C for 10 min. After thermal cycling, the droplets were read and analyzed by QX200 Droplet Reader (BIO-RAD). The detail of workflow including droplet generation, droplet transfer, heat sealing was conducted according to the manufacturer’s protocol.

### Microfluidic chip design and fabrication

The overall architecture of our microfluidic chip simply consists of a single-piece of polydimethylsiloxane (PDMS) irreversibly bonded on a slide glass (**Supplementary Fig.1a-b**). The microfluidic chip has one sample inlet for the dispersed phase, a pair of oil inlets for the continuous phase, and an outlet connecting to a syringe. These inlets open to the atmosphere and are maintained at an almost constant pressure (~1 atm), a pressure differential through the chip is created to actuate the fluids when the piston of the connecting syringe is pulled outward and locked in a setting volume. A flow-focusing geometry is adopted for droplet generation, followed by a Christmas tree-shaped distribution channel network. This design facilitates constructing high-density monolayer droplet array within the fluorescence imaging chamber.

### Droplet generation

Prior to droplet generation, the microfluidic chip is filled with oil phase (light mineral oil containing the stabilizing surfactants 3% ABIL EM 90 (Degussa) and 0.1% w/w Triton X-100. Then, a single-use plastic syringe is connected to the out inlets of the chip with a length of Teflon tubing (0.6 mm×0.9 mm, [i.d.× o.d.]). When the capillary force-driven oil flows reach to the sample inlet, a 3-μl aliquot of Cas13a mix loaded into the sample inlet with a micropipette, ensuring no air-bubble to be trapped into this inlet. To initiate the droplet generation, the piston of the syringe is pulled outward, and locked in place with one-milliliter volume using a binder clip. After the imaging chamber is filled with the microdroplets, the piston is gently pushed back its starting position, and then the tubing is cut off.

### Droplet Imaging

The prepared microdroplets along with the microchip is underwent a 60-min isothermal reaction using a heating plate. Subsequently, the microfluidic chip is continuously imaged with an Olympus IX-71 inverted microscope to record the bright field and fluorescence images of the microdroplets. A 10× magnification objective lens is capable of imaging 1045 ×1390 μm^2^ areas, allowing almost 1200-2400 (with different droplet density) microdroplets to be captured in a single snapshot. For a single test, the bright field images are used for accumulating the total number of the microdroplets (at least 20,000 droplets), whereas the corresponding fluorescence images (ISO 1600, 4-s exposure time) are used to discriminate the positive droplets with a signal-to-noise ratio above 3. To determinate the average size of the microdroplets, three randomly chosen areas were imaged with a 40× objective and over 30 droplets in these microscopic images were measured with cellSens^®^ software (Olympus).

### Digital quantification

According to the Poisson distribution, the measured concentration (*C*) was calculated by

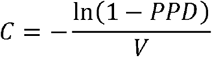

Where droplet volume (*V*) was estimated according to the diameter (*D*),

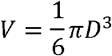

and *PPD* was the proportion of positive droplets.

## Acknowledgments

We thank Professor Yanli Wang for providing pET28a-His6-SUMO-LbuCas13a plasmids for expressing LbuCas13a. We also thank Zhenguo Zhu for helpful software programing in droplet counting. This work is supported by the National Natural Science Foundation of China (21804044, 21475048, and 21874049), Guangzhou Science and Technology Program (201904010413), the Fundamental Research Funds for the Central Universities (D2192220), the National Science Fund for Distinguished Young Scholars of Guangdong Province (2014A030306008), and the Special Support Program of Guangdong Province (2014TQ01R599).

## AUTHOR CONTRIBUTIONS

X.M.Z., T.T. and B.W.S. conceived the concept and designed the study. T.T. and X.M.Z. designed and constructed the Cas13 system. T.T., and L.L. carried out cell culture, total small RNA extraction, bulk fluorescence detection, ddCA, qRT-PCR and ddPCR experiments. B.W.S. designed and constructed microfluidic droplet chip and analysis platform for ddCA. B.W.S., T.T., and X.M.Z. interpreted the results and wrote the manuscript. All authors commented on and edited the manuscript.

