## Supplemental Information for "Droplet-digital Cas13a assay enables direct single-molecule microRNA quantification"

^1^College of Biophotonics & School of Life Science, South China Normal University, Guangzhou 510631, China, ^2^Department of Laboratory Medicine, Guangzhou First People’s Hospital, the Second Affiliated Hospital of South China University of Technology, Guangzhou 510180, China, ^3^Clinical Molecular Medicine and Molecular Diagnosis Key Laboratory of Guangdong Province, Guangzhou 510180, China, ^4^Institute for Brain Research and Rehabilitation, Guangdong Key Laboratory of Mental Health and Cognitive Science, Center for Studies of Psychological Application, South China Normal University, Guangzhou 510631, China,

**Contents**

**Supplementary Note 1.** Comparison of the current methods for microRNA quantification.

**Supplementary Note 2.** DNA and RNA sequences used in this study.

**Supplementary Note 3.** Supporting discussion.

**Supplementary Figures**

**Fig.S1** Photograph of the assembled microfluidic droplet chip.

**Fig.S2.** Microscopic photography of droplet array.

**Fig.S3.** Real-time fluorescence measurement of miRNA in bulk reaction volume based on Cas13a system.

**Fig.S4.** Detection of four microRNAs from different families using ddCA.

**Fig.S5.** Bulk fluorescence detection of microRNA members within a family.

**Fig.S6.** Characterization of LbuCas13a purification.

**Fig.S7.** Characterization of LbuCas13a crRNAs produced by in vitro transcription.

**Supplementary Video 1.** High throughput droplet generation using a microfluidic droplet chip.

**Supplementary References**

**Supplementary Note 1. Comparison of the current methods for microRNA quantification**

|  | Sensitivity | Specificity | Dynamic range | Absolute quantification | Amplification free | Manipulations | Time to result | Infrastructure  requirement |
| --- | --- | --- | --- | --- | --- | --- | --- | --- |
| qRT-PCR^1-3^ | zM to fM | medium | 7-logs | 🗙 | 🗙 | RT^(e)^ | hours | few |
| ddPCR^4,5^ | single molecule | medium | 5-logs | ✔ | 🗙 | RT | hours | moderate |
| Microarray^3,6,7^ | nM to mM | low | 4-logs | 🗙 | ✔ | labelling | days | moderate |
| RNA-Sequencing^3,8,9^ | zM to fM | high | ≥5-logs | 🗙 | 🗙 | RT and ligation | weeks | substantial |
| FCS^(a)10^ | fM to pM | high | 3-logs | ✔ | ✔ | none | hours | moderate |
| DNA nanomechanics^11^ | aM to fM | low | 3-logs | ✔ | ✔ | probe immobilization | hours | moderate |
| Nanopore-based detection^12,13^ | fM to pM | high | 4-logs | ✔ | ✔ | none | hours | few |
| SiMREPS^(b)14^ | fM to pM | high | 2 or 3-logs | ✔ | ✔ | probe immobilization | hours | moderate |
| Simoa^(c)15^ | aM to fM | high | 4-logs | ✔ | ✔ | washing | hours | moderate |
| Nanoparticles^16^ | aM to fM | medium | 5-logs | 🗙 | ✔ | washing and centrifugation | hours | few |
| miracles^(d)17^ | fM to pM | high | 6-logs | 🗙 | ✔ | none | hours | few |
| ddCA(this work) | single molecule | high | 5-logs | ✔ | ✔ | none | hours | few |

(a)FCS= fluorescence correlation spectroscopy. (b)SiMREPS= single-molecule recognition through equilibrium passion sampling. (c) Simoa=single molecule array. (d) miracles= microRNA-activated conditional looping of engineered switches. (e) RT= reverse transcription.

**Supplementary Note 2.** DNA and RNA sequences used in this study.

| Name | Sequence (5’-3’) |
| --- | --- |
| miR-17 | CAAAGUGCUUACAGUGCAGGUAG |
| miR-106a | AAAAGUGCUUACAGUGCAGGUAG |
| miR-20a | UAAAGUGCUUAUAGUGCAGGUAG |
| miR-20b | CAAAGUGCUCAUAGUGCAGGUAG |
| miR-10b | UACCCUGUAGAACCGAAUUUGUG |
| miR-21 | UAGCUUAUCAGACUGAUGUUGA |
| miR-155 | UUAAUGCUAAUCGUGAUAGGGGU |
| FQ5U | FAM-UUUUU-BHQ1 |
| crRNA- miR-17 | GACCACCCCAAAAAUGAAGGGGACUAAAACCCUGCACUGUAAGCACUUUG |
| crRNA- miR-17-M | GACCACCCCAAAAAUGAAGGGGACUAAAACCCUGCACUGUAAGCAGUUUG |
| crRNA- miR-106a-M | GACCACCCCAAAAAUGAAGGGGACUAAAACCCUGCACUGUAAGCAGUUUU |
| crRNA- miR-20a-M | GACCACCCCAAAAAUGAAGGGGACUAAAACCCUGCACUAUAAGCAGUUUA |
| crRNA- miR-20b-M | GACCACCCCAAAAAUGAAGGGGACUAAAACCCUGCACUAUGAGCAGUUUG |
| miR-17-stem-loop-RT primer | GTCGTATCCAGTGCAGGGTCCGAGGTATTCGCACTGGATACGACCTACCT |
| miR-17-Forward-PCR primer | GCGGCGCAAAGTGCTTAC |
| miR-17-Reverse-PCR primer | CCAGTGCAGGGTCCGAGGTA |
| miR-17-PCR-fluorescence probe | FAM-TGGATACGACCTACCTG -MGB |
| T7 promoter | TAATACGACTCACTATAGG |

**Supplementary Note 3.** **Concept for single-molecule miRNA quantification using ddCA**

The reaction volume might be a key issue for Cas13a-based single-molecule miRNA detection. Assuming that one target miRNA molecule recognized by Cas13a could induce the activation of quenched fluorescent reporter with a turnover of N, and a fluorescence detector requires a critical concentration of activated fluorescent reporter up to C* due to its detection limit, it requires at least (V·C*/N) target molecules to reach this concentration in a detection volume of V. Consequently, if the detection volume is reduced to a critical volume of V*=N/C*, the single-molecule miRNA detection is achieved. Another reason for such superior performance is derived from droplet-powered digital assay. The droplet digital quantitation converts the continuous measurement of an amplitude of a physical quantity (e.g., fluorescence intensity or initial velocity) in bulk volume to a calculation of binary end-point signals adopting either a positive or negative value if the droplet contains at least one target molecule or none, thus enabling to provide precise and absolute miRNA quantification without the need for internal reference or calibration.

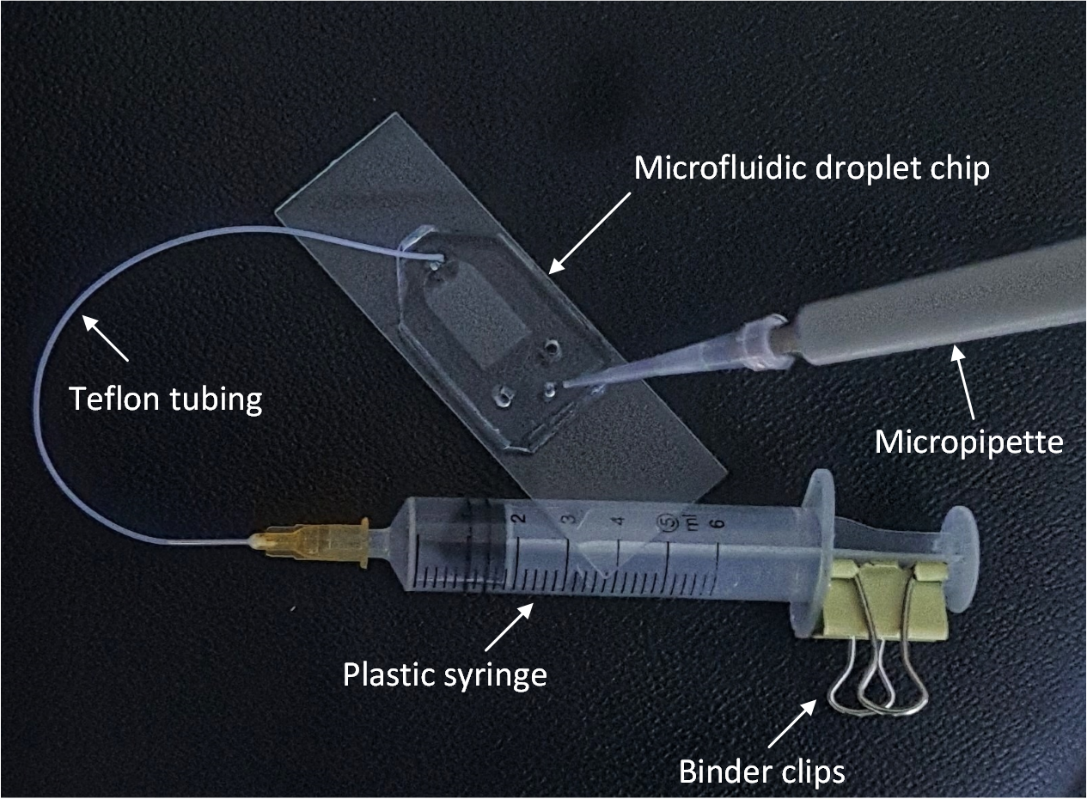

**Figure S1.** Photograph of the assembled microfluidic droplet chip with all necessary equipment for sample loading and droplet generation.

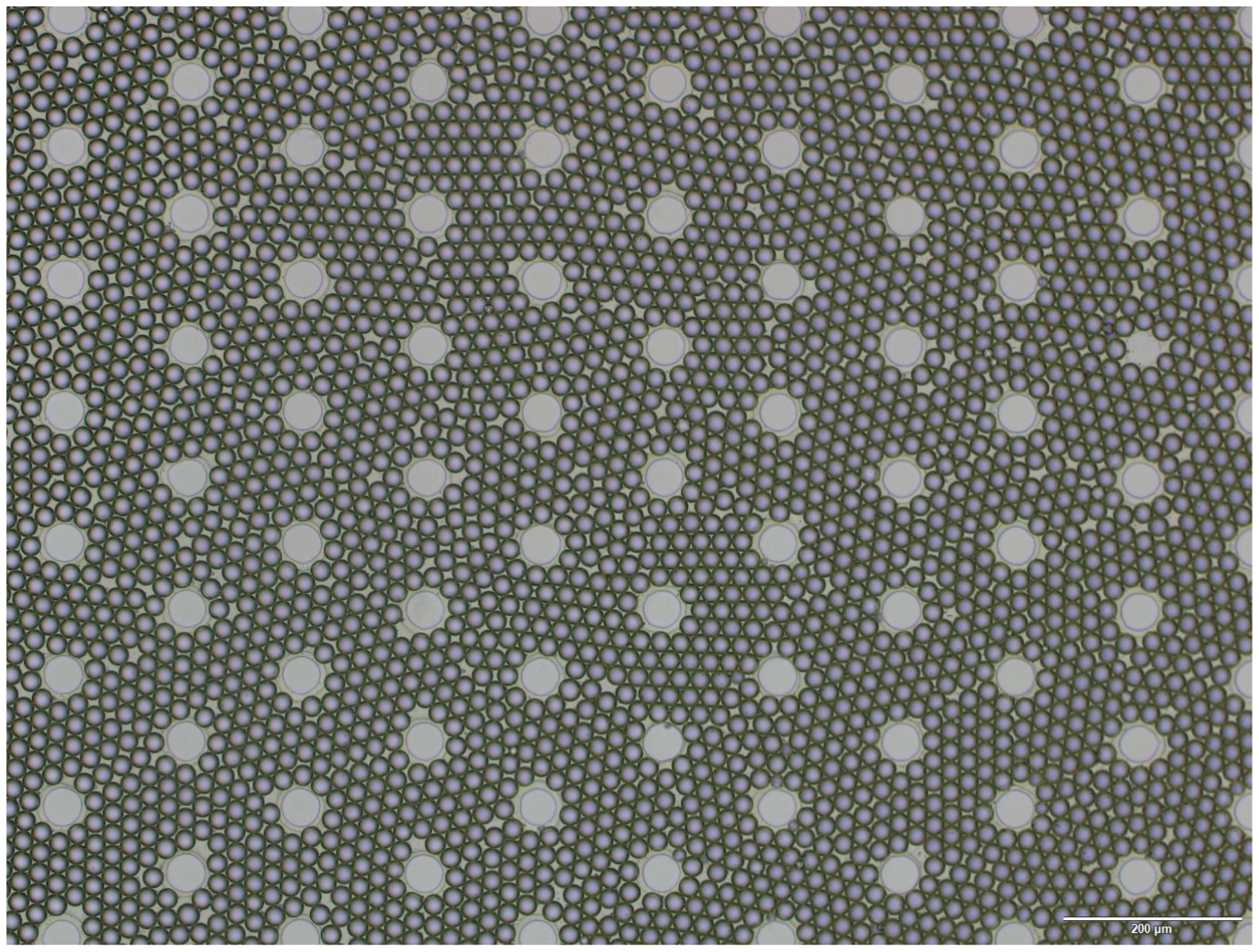

**Figure S2. Microscopic photography of the self-assembled droplet array.** Microscopic photography of the droplets under 10 x objective lens (scale bar: 200 μm).

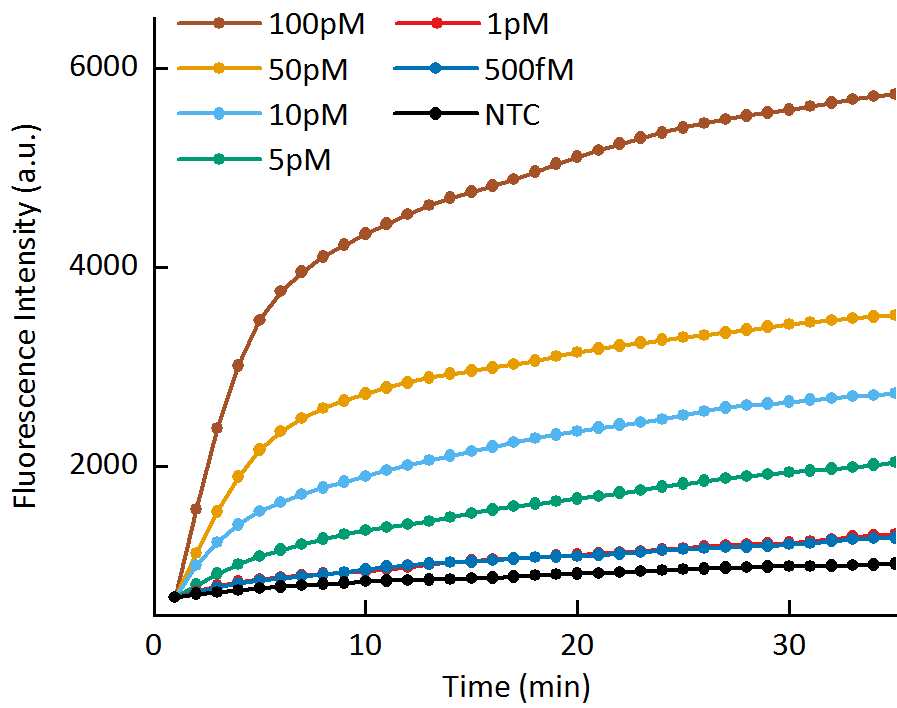

**Figure S3.** Real-time fluorescence measurement for miR-17 detection in bulk reaction volume (25 μL) using fluorescence detection system for 35 minutes, LOD= 500 fM.

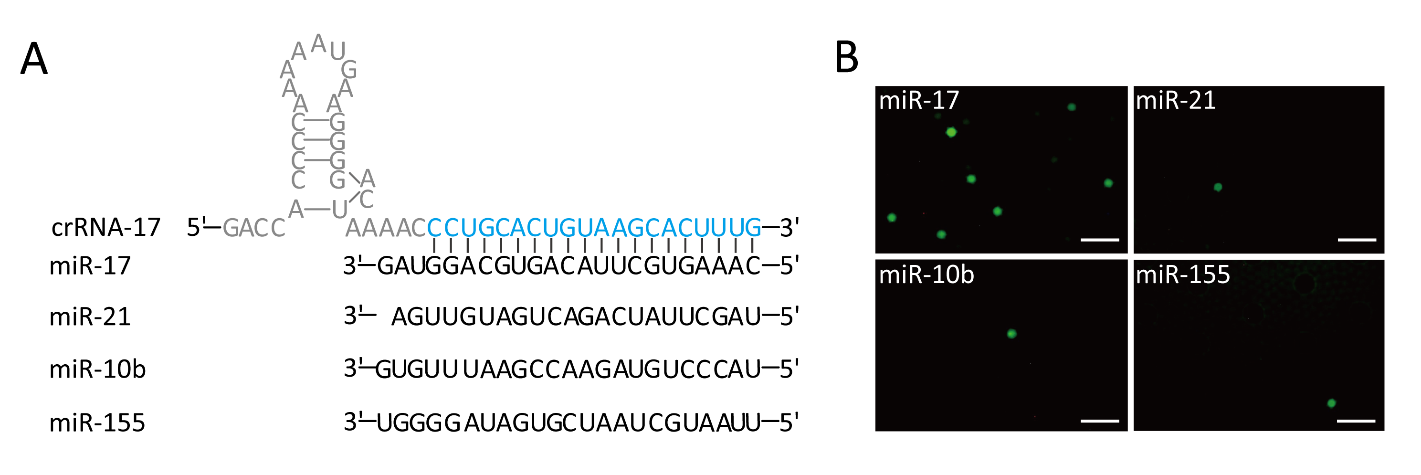

**Figure S4. Detection of four microRNAs from different families using ddCA.** (A) Sequences of crRNA with a 20-nt spacer (blue) complementary to miR-17, and three microRNAs from other families. (B) Representative microscopy images of detecting four microRNAs using the same input target concentration (10 fM) (scale bar: 100 μm).

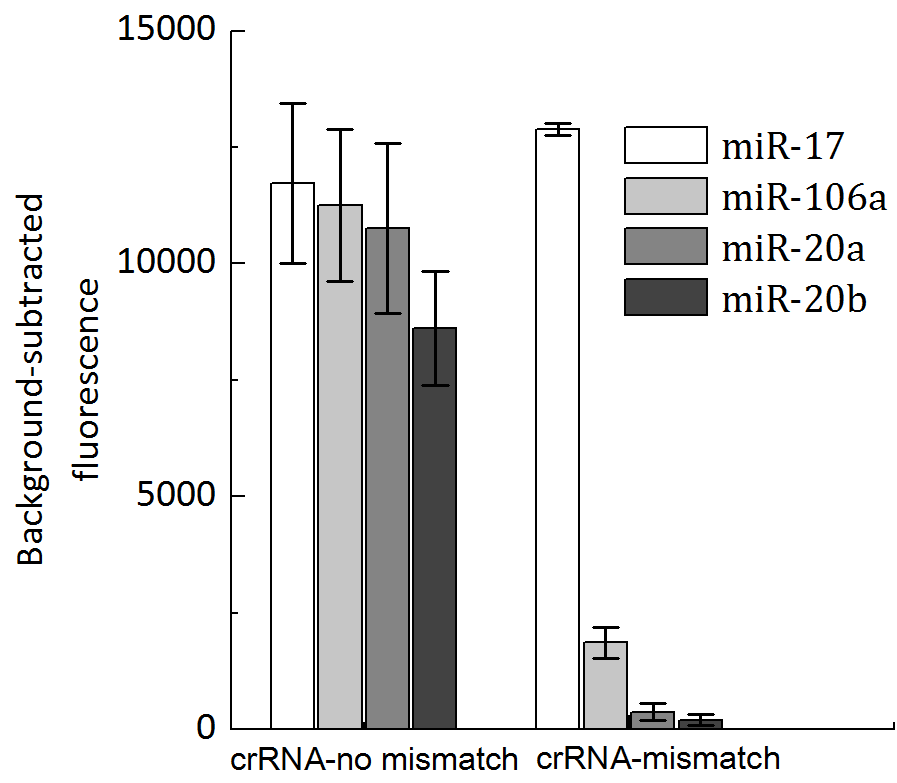

**Figure S5. Bulk fluorescence detection of miR-17 family used complementary miR-17 crRNA or one base artificially mutated miR-17 crRNA.** Synthetic mismatched crRNA provides more robust discrimination than non-mismatch crRNA. (n=3 technical replicates, bars represent mean ± s.e.m.)

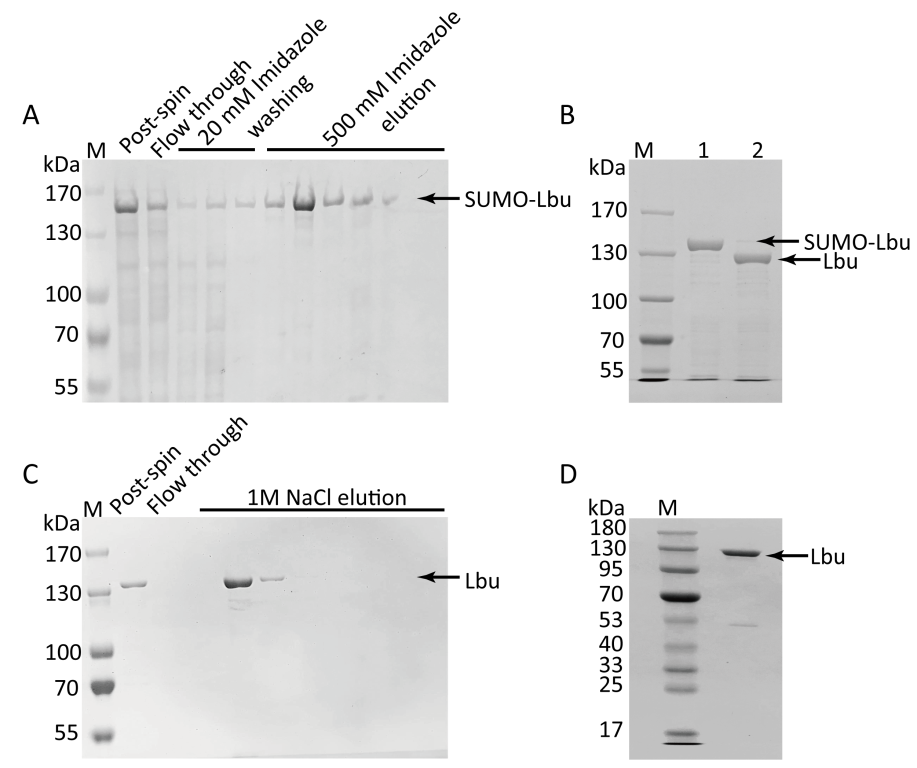

**Figure S6. Characterization of LbuCas13a purification.** (**A**) Coomassie blue stained SDS-PAGE gel of purified SUMO-LbuCas13a stepwise eluted from Ni-NTA column. (**B**) SUMO-LbuCas13a before (lane1) and after (lane2) digestion. SUMO tag was removed by Ulp. (**C**) Coomassie blue stained SDS-PAGE gel of purified LbuCas13a stepwise eluted from Heparin column. (**D**) Coomassie blue stained SDS-PAGE gel of concentrated LbuCas13a.

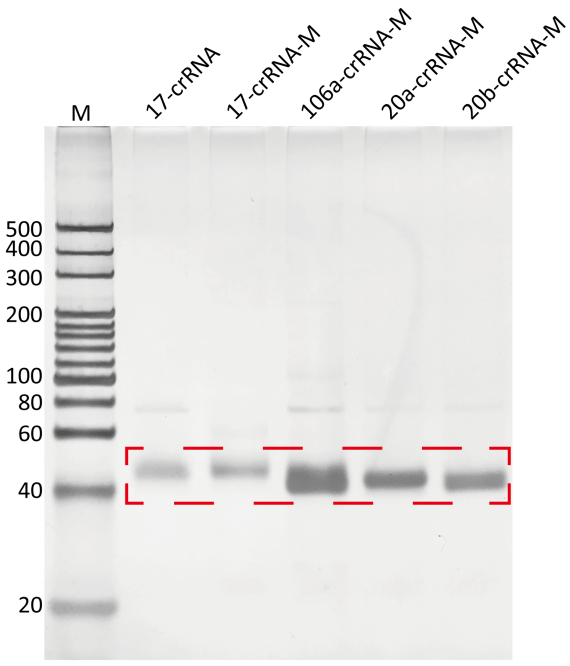

**Figure S7. Characterization of LbuCas13a crRNAs produced by *in vitro* transcription.**
